# Tropical Soil Metagenome Library Reveals Complex Microbial Assemblage

**DOI:** 10.1101/018895

**Authors:** Kok Gan Chan, Zahidah Ismail

## Abstract

In this work, we characterized the metagenome of a Malaysian mangrove soil sample via next generation sequencing (NGS). Shotgun NGS data analysis revealed high diversity of microbes from Bacteria and Archaea domains. The metabolic potential of the metagenome was reconstructed using the NGS data and the SEED classification in MEGAN shows abundance of virulence factor genes, implying that mangrove soil is potential reservoirs of pathogens.

## Introduction

Mangrove forests are usually located at the tropical and subtropical latitudes. They are present at the transition of land and sea which makes them susceptible to tidal change and salinity. Mangrove soils generally comprise soft, muddy and anaerobic sediment with thin top layer of aerobic sediment. They also function as heavy metal sink^1^ which acts as natural sink and filtration system. Mangrove swamps are the habitat for a diverse variety of fauna especially juvenile fishes, and they also act as breeding and nursery grounds for these aquatic animals.

Microorganisms in the mangrove habitat play an important role in maintaining the productivity, conservation and nutrients of this ecosystem. Microorganisms are involved in biogeochemical cycles that supply nutrients to plants and animals^2, 3^. Mangroves are rich in organic matters but usually lack phosphorus and nitrogen^4-7^. Their microorganisms’ activities are high because they are very efficient in recycling the nutrients contained therein. Microorganisms are directly involved in nitrogen fixation, phosphate solubility, photosynthesis, sulfate reduction and production of other substances.

The mangrove environment is highly susceptible to anthropogenic effects such as pollution, deforestation and human activity. These could change the dynamic mangrove ecosystem which in turn affects the mangrove community and disturbs the microorganism community that maintains the productivity and conservation of mangrove.

This study aimed to investigate the metagenome of a mangrove soil sample and their ecological role via metabolic reconstruction. We used the Illumina HiSeq 2000 platform to carry out shotgun metagenome next generation sequencing (NGS). This method avoided bias of PCR amplification as in the case of amplicon sequencing and it enabled parallel study on both the taxonomic and functional diversities. We hypothesized that the abundance and diversity of microbes and their functional attributes to be similar to those of previous studies^8, 9^.

## Materials and Methods

Sampling was done on a soil sample obtained in the east coast of Peninsular Malaysia, namely Rantau Abang (RA) (N04° 54.189’ E103° 22.208’). No specific permissions were required for the chosen locations and such research activities. Our work also did not involve endangered or protected species. The top 5 to 20 cm of soil was collected and stored at -20°C until processing. A portion of the soil sample was sent for biochemical analyses of its pH, carbon nitrogen ratio, and contents of phosphorus, sulfur and heavy metals, like arsenic, cadmium, lead and mercury, as described previously^10^.

DNA extraction was carried out according to the protocol as described previously^11^ with modifications. Traces of plant materials were removed from the soil prior to extraction. Briefly, 5g of soil was added with 13.5ml of DNA extraction buffer (Tris-HCl, pH8 100mM; EDTA, pH8 100mM; Na_2_HPO_4_, pH7.8 100mM; NaCl, 1.5M; CTAB, 1% w/v), 100μl of proteinase K (10mg/μl), and 200μl of lysozyme (10mg/μl). The mixture was incubated horizontally at 37°C with orbital shaking (225rpm). After 30min, 0.5ml of SDS (20% w/v) was added and the mixture further incubated in a 65°C water bath for 2h with gentle mixing by inverting the tube at 15min intervals. The supernatant was collected by centrifugation at 6000 × *g* for 10min. The pellet was suspended in 4.5ml of DNA extraction buffer and 0.5ml of SDS (20% w/v) and vortexed for 10s followed by incubation at 65°C for 10min. The supernatant was then collected by centrifugation and pooled with the supernatant collected previously. Equal volume of chloroform:isoamyl alcohol (24:1, vol/vol) was added to the pooled supernatant and the mixture was gently mixed by inversion. The aqueous phase was transferred to a clean, sterile tube after centrifugation at 6000 × *g* for 10min. The chloroform:isoamyl alcohol step was repeated once. For DNA precipitation, 0.6 volume of cold isopropanol was added and the resultant mixture incubated at -20°C for 30min. DNA was collected by centrifugation at 16000 × *g* for 20min, followed by washing with 70% (v/v) ethanol and kept in -20°C for 15min. Ethanol was removed by centrifugation at top speed in a table top centrifuge for 10min and the pellet was air dried aseptically. The DNA pellet was then dissolved in elution buffer (Roche High Pure PCR Product Purification Kit).

The soil metagenomic DNA was further purified by gel elution in a 3% (w/v) low melting temperature agarose electrophoresis. The metagenomic DNA was mixed with 80% (v/v) glycerol and 6× loading dye and the mixture was then loaded into a well. Electrophoresis was carried out at 15V for 16 to 20h. DNA was excised from the gel with a sterile blade and recovered using the Qiagen Gel Extraction Kit (Venlo, Netherlands). DNA concentration and purity were determined using Qubit and Nanodrop 2000c by Life Technologies, respectively. The purified DNA was then subject to NGS using Illumina HiSeq 2000.

For taxonomic analysis, the metagenomic nucleotide sequences obtained were trimmed using CLC Bio Genomic Workbench 5.5.2 (Aarhus, Denmark) at 50-nucleotide length to remove short low quality reads. The trimmed data were then blasted against the NCBI Microbial database (dated 22 Jan 2013) using Blastall 2.2.25 (NCBI) at the expected value of 1×10^-20^.

For functional gene study, the trimmed nucleotide sequences were assembled using *de novo* assembly in CLC Bio Genomic Workbench at the minimum contig length of 400 nucleotides. The assembled data were extracted at coverage of ≥10%. Gene prediction was performed on the extracted sequences using Prodigal 2.60^12^, and each predicted gene was annotated using RAPsearch 2.09^13, 14^ (against the NCBI NR database (dated 22 January 2013). Data obtained for both taxonomic and functional distributions were analyzed in MEGAN 4.70.4^15^. Taxonomic analysis was done according to the percentage identity filter to get the best sequence match. In MEGAN, functional analysis was accomplished with the SEED^16^ classification.

The data of this study are available as NCBI database accession number SRR748204.

## Results

### Biochemical Analyses

Table 1 shows the biochemical properties of the Rantau Abang (RA) soil sample. The pH for RA sample was recorded as pH of 5.1.

**Table 1.** Results of biochemical analyses of the soil sample. ND: not detected.

| Parameter | Unit | RA |
| --- | --- | --- |
| Arsenic (As) | mg/kg | ND(<0.5) |
| Cadmium (Cd) | mg/kg | 0.10 |
| Lead (Pb) | mg/kg | 25.34 |
| Mercury (Hg) | mg/kg | ND(<0.05) |
| pH (10% w/w) | - | 5.1 |
| Phosphorus | mg/kg | 449.70 |
| Sulfur | mg/kg | 1374.52 |
| Carbon:Nitrogen | % | 5.19:0.31 |

### Metagenomic Library Analysis

The RA sample metagenome library shows very high nucleotides sequenced after editing and >500,000 contigs generated at the coverage at approximately 41 times (Table 2).

**Table 2.**
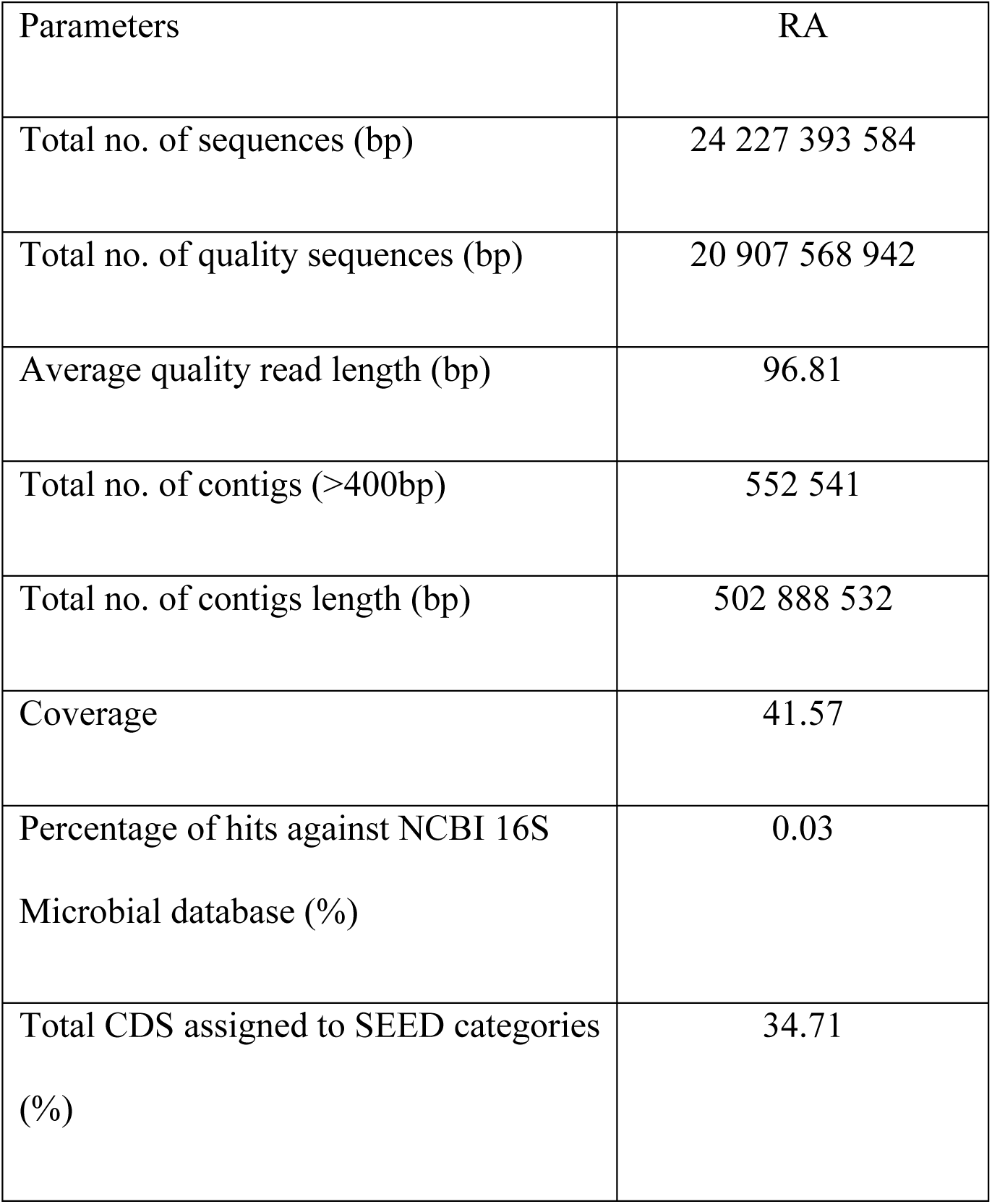
Reads statistics. The numbers of reads were generated by Illumina HiSeq 2000. The RA sample showed good quality of reads in term of number, length and number of contigs generated.

### Microbial Taxonomic Distribution

A total of 98% of the reads from RA sample was assigned to the domains level by MEGAN and they excluded the “No hits” reads category. The majority of the assigned reads from RA samples was of the domain Bacteria while the remaining were of the domain Archaea with 78.52% reads assigned to Bacteria and 21.48% to Archaea (Figure 1).

**Figure 1.**
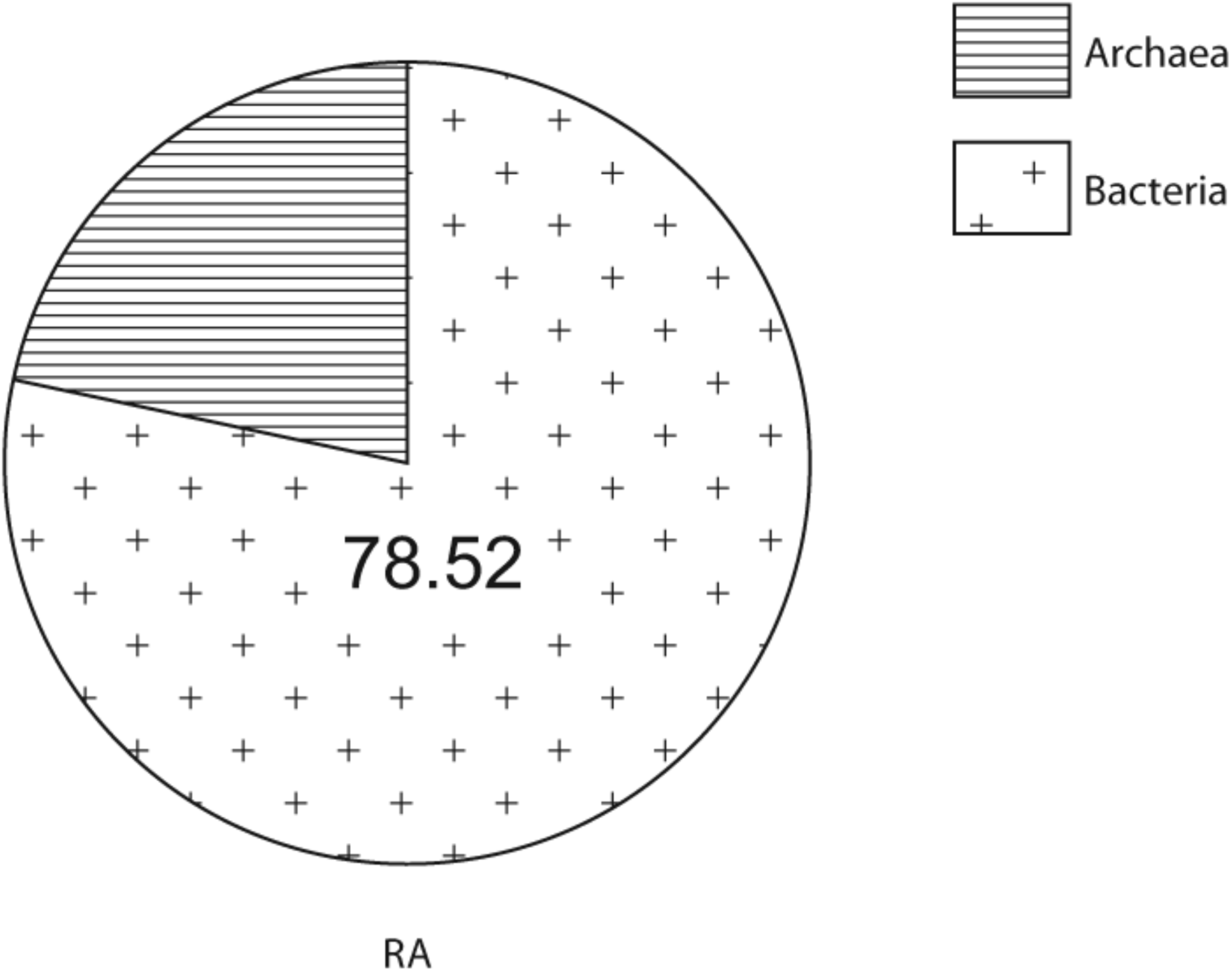
Bacteria and Archaea. The majority of the reads were assigned to the domain Bacteria by MEGAN. The number of reads detected for archea was significantly higher than previously reported.

There were 27 phyla hits from the domain Bacteria for RA metagenome library. The phylum Proteobacteria dominated other phyla in (43.72%) in the RA sample. There were 10 phyla with abundance percentages of more than 1% in the RA sample. In this sample, the phyla detected were Proteobacteria, Acidobacteria (17.68%), Firmicutes (13.45), Actinobacteria (4.55%), Nitrospirae (4.22%), Planctomycetes (3.06%), Chloroflexi (2.88%), Verrucomicrobia (2.69%), Spirochaetes (1.70%), Chlamydiae (1.32%) and Bacteroidetes (1.31%) (Figure 2a). In RA sample, unclassified bacteria were clustered as Caldithrix, Haloplasmatales and some phototrophic bacteria.

**Figure 2a.**
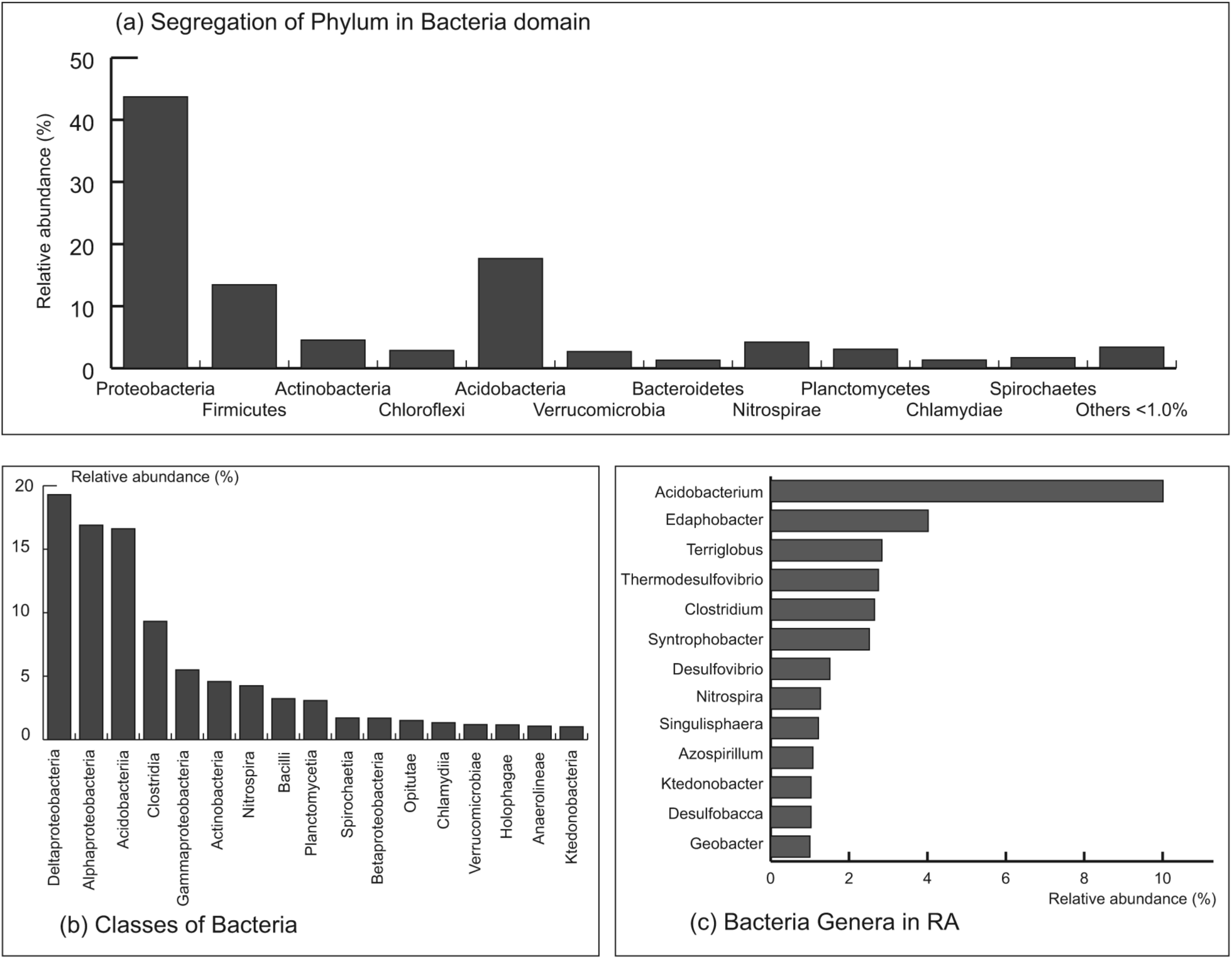
The segregation of phyla in the domain Bacteria. Proteobacteria was the dominant phylum in the samples with almost half of the reads assigned to this particular phylum. **Figure 2b Classes of bacteria.** Classes of bacteria that were present in the RA sample at more than 1%. **Figure 2c Bacterial genera.** The genera shown are those that had more than 1% reads. Bacterial genera detected in RA sample in which *Acidobacterium* was the most abundant genus.

Forty-three classes of bacteria were detected in RA sample (Figure 2b) where the first two most abundant classes were Deltaproteobacteria (19.29%) and Alphaproteobacteria (16.89%), followed by Acidobacteria (16.61%) (Figure 3b). The other minor classes included Clostridia (9.32%), Gammaproteobacteria (5.50%), Actinobacteria (4.58%), Nitrospira (4.24%), Bacilli (3.24%), Planctomycetia (3.08%), Spirochaetia (1.71%), Betaproteobacteria (1.69%), Opitutae (1.52%), Chlamydia (1.33%), Verrucomicrobiae (1.19%), Holophagae (1.17%), Anaerolineae (1.06%), and Ktedonobacteria (1.02%).

At the genus level, *Acidobacterium* of the Acidobacteria phylum was the dominant genus in RA sample. The abundance frequencies of this genus is 10.01% (Figures 2c).

Five classes of Proteobacteria namely Alphaproteobacteria, Betaproteobacteria, Deltaproteobacteria, Gammaproteobacteria and Epsilonproteobacteria, were detected in RA sample). Among these, Deltaproteobacteria (43.88%) was the major class, followed by Alphaproteobacteria (38.43%), Gammaproteobacteria (12.51%) Betaproteobacteria (RA 3.85%) and Epsilonproteobacteria (RA 1.33%) (Figure 3). The segregation of orders within Deltaproteobacteria (Figure 4) showed Syntrophobacterales was the most abundant order in this soil sample. At the genus level of this order *Syntrophobacter* was the most abundant genus (Figures 5).

**Figure 3.**
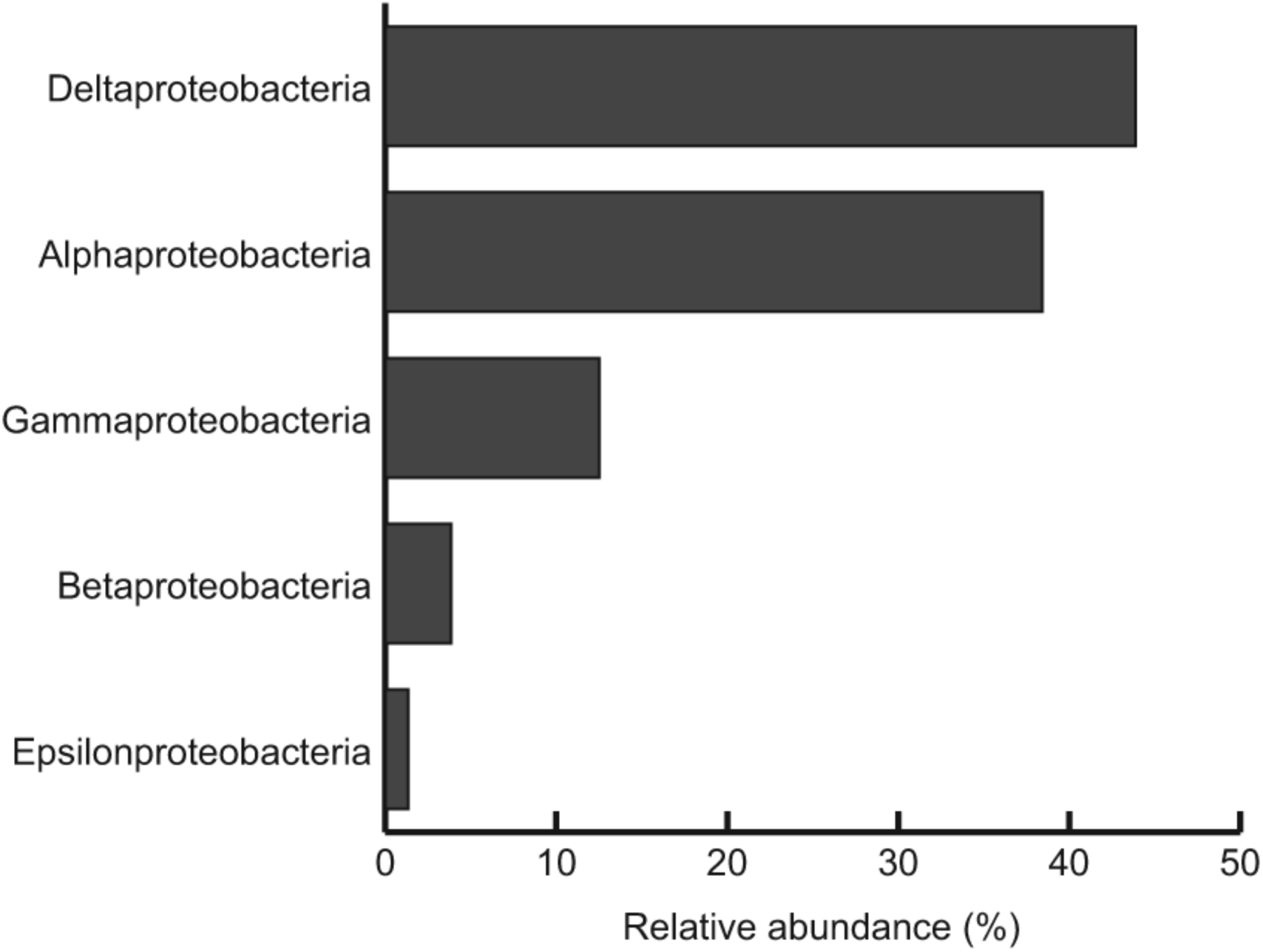
Classes of Proteobacteria. Deltaproteobacteria was the most abundant class in RA samples.

**Figure 4.**
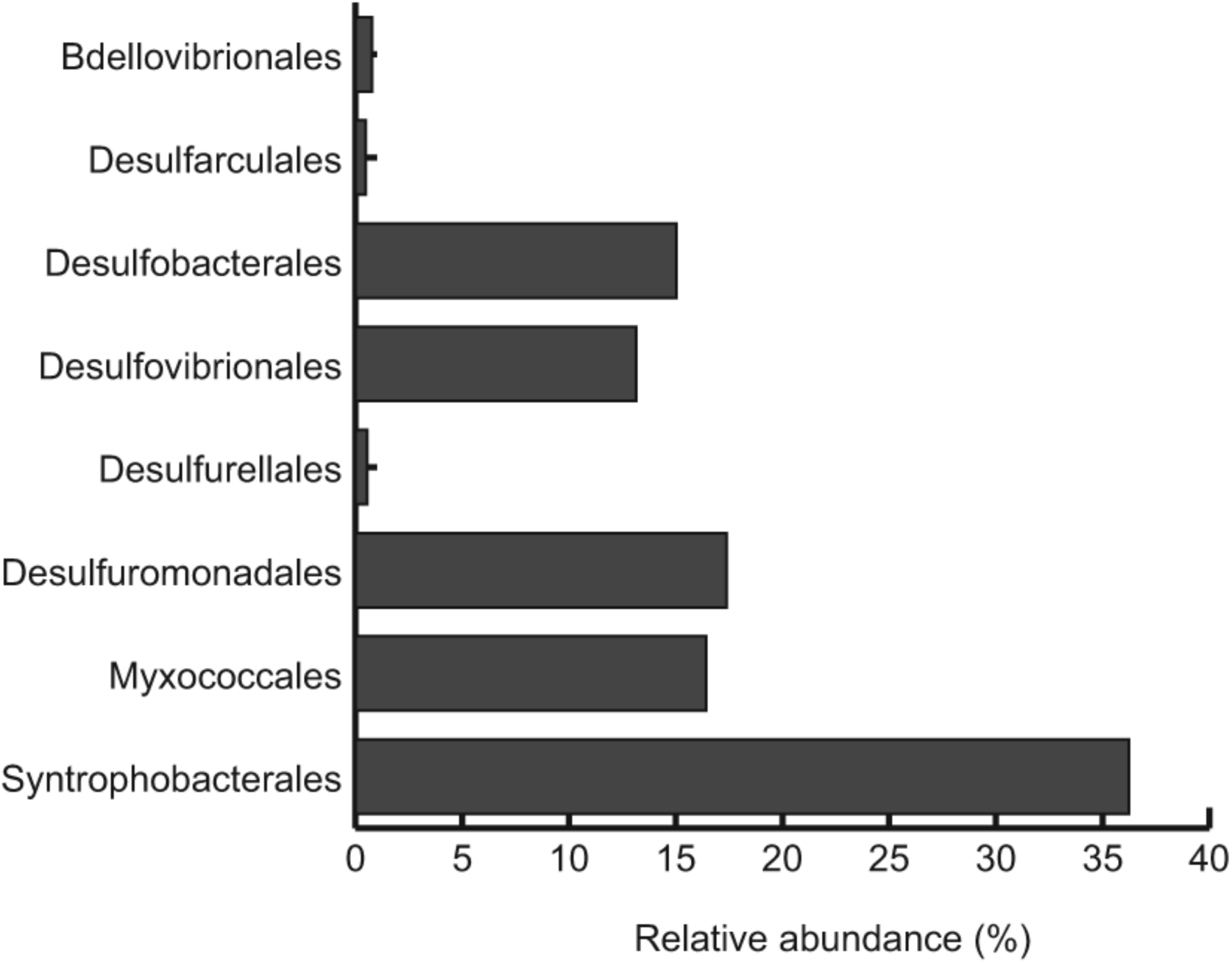
Order level of Deltaproteobacteria. Syntrophobacterales were the most abundant order.

**Figure 5.**
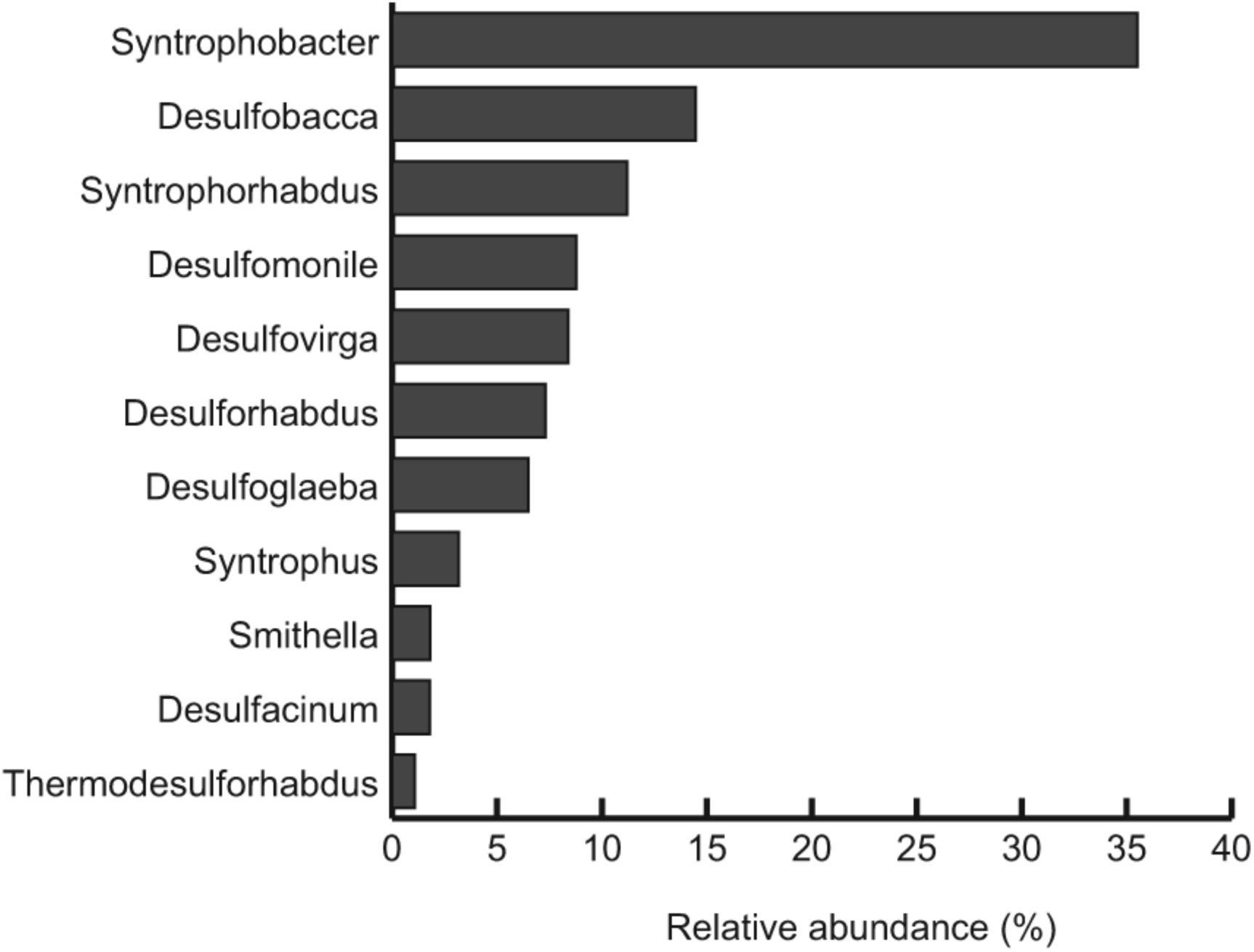
The most abundant genera in Syntrophobacterales. *Syntrophobacter* was the most abundant genus.

The RA sample showed present of archaea but only phyla Crenarchaeota and Euyarchaeota were detected. In the RA sample, Crenarchaeota (63.78%) was present at a higher percentage as compared to Euyarchaeota (36.22%) (Figure 6). A total of eight classes of archaea were detected in both soil samples and among them, Thermoprotei (RA 63.78%) and Methanomicrobia (RA 17.85%) were the two dominant classes. Other minor archaea classes present in the RA sample were Thermococci (5.48%), Methanococci (5.35%), Thermoplasmata (3.52%), Methanobacteria (2.17%), Archaeoglobi (1.36%) and Halobacteria (0.47%).

**Figure 6.**
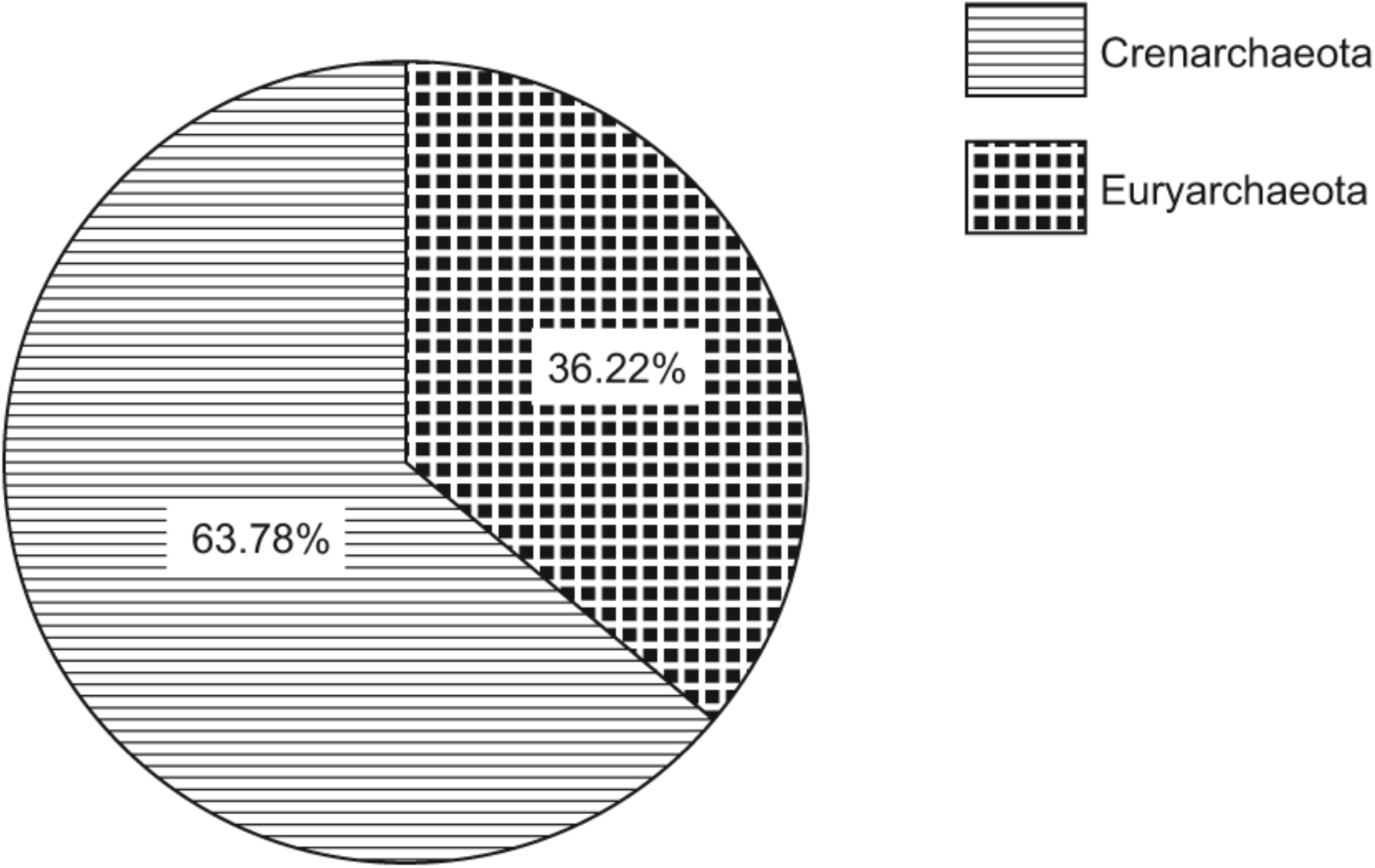
Distribution of archaeal phyla in the sample. RA sample possessed high percentage of Crenarchaeota.

The taxonomic diversity for the domains Bacteria and Archaea was estimated at the genus level using the Shannon-Weaver diversity index, H’, in MEGAN and the H’ value RA samples was 7.765.

### Metabolic Functional Analysis via Reconstruction of Metagenome Library

The gene anthology was derived from the SEED classifications. Using this approach, the most abundant gene detected in the RA sample was associated with carbohydrate metabolism (12.97%). The second most abundant genes in the RA sample were associated with protein metabolism (9.89%), virulence (9.53%), respiration (8.39%) and amino acids and their derivatives (8.19%) (Figure 7).

**Figure 7.**
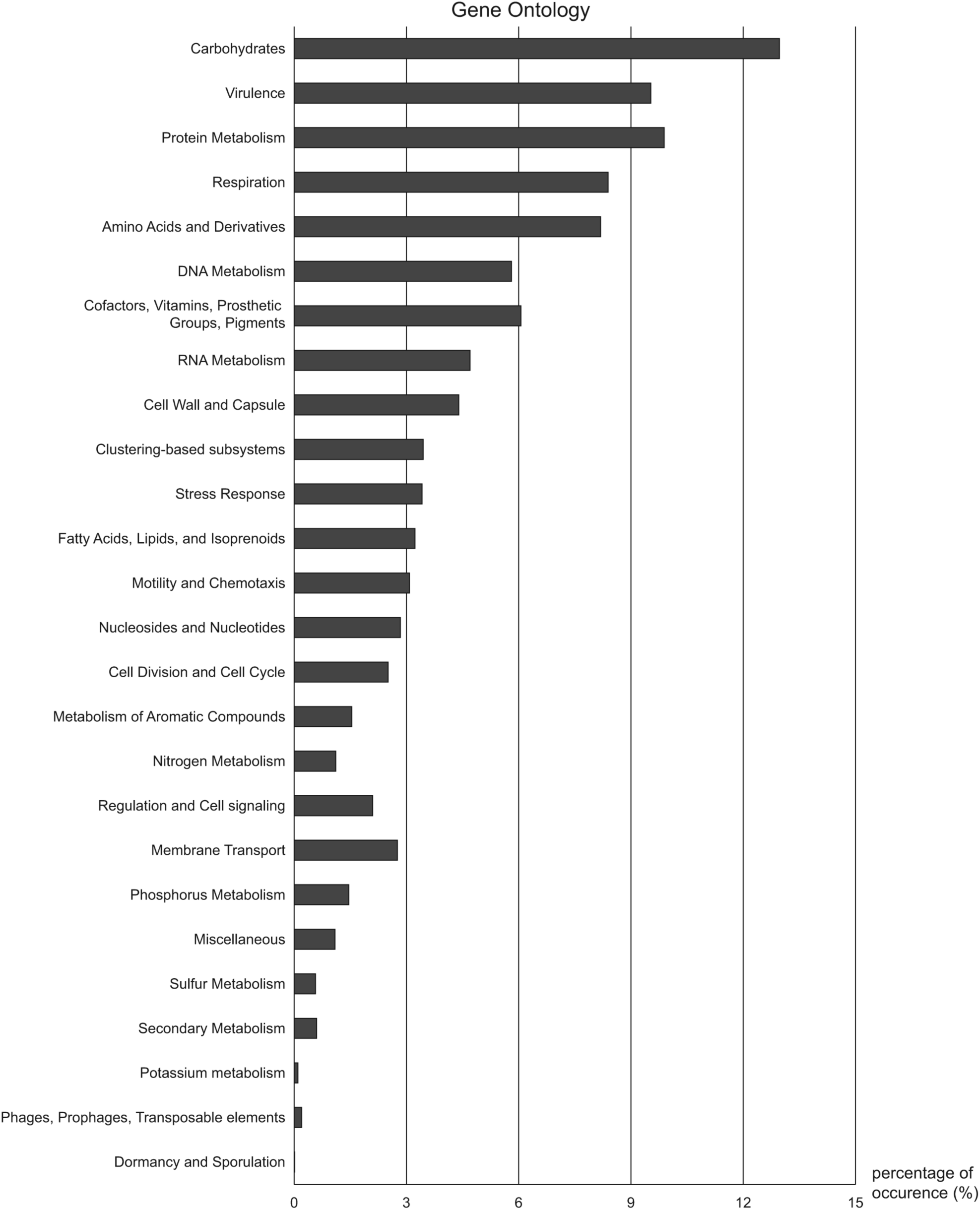
Gene ontology. The SEED classification by MEGAN.

## Discussion

The application of the NGS enables the study of microbial diversity and function in metagenomes without the need of culturing bacteria thus bypassing the growing of fastidious bacteria, which are often unculturable, on laboratory media. However, this method depends on the reliability of NGS data generated.

Most bacteria detected in the RA metagenomes were either anaerobic or facultative anaerobic. They were predominantly from the domain Bacteria. However, in the RA metagenome, a significant number of bacteria belonging to the domain Archaea were detected and the percentage of archaeal abundance (21.48%) is significantly higher than the percentage values reported for other soil metagenomes^17^. Crenarchaeota is known to be present at high sulfur content environment and for its ability to utilize sulfur^18^. The high percentage of this phylum detected in the RA sample is consistent with the high concentration of sulfur in the RA sample.

The bacterial diversity detected in the RA sample conforms to the common soil bacteria present in other types of soil in other geographical locations^19^. In our RA sample, Proteobacteria remains the most abundant bacteria and it comprised five different classes. However, the distribution of Proteobacteria classes in this study differs from the distributions of Proteobacteria classes reported for other mangrove habitats^8, 20^. In contrast to other mangrove metagenomes reported to date, both the Malaysian mangrove metagenome possessed Deltaproteobacteria as the dominant class of Proteobacteria. Even though Proteobacteria was the dominant phylum in RA metagenome, *Acidobacterium* of the Acidobacteria phylum was the most abundant genus in our soil sample.

The presence of the high frequency of genes associated with carbohydrate metabolism in our soil metagenome analysis is not surprising because these genes are commonly detected in abundance in most studies of soil metagenomes^8, 21^. However, the presence of the high frequency of virulence factor genes in our soil metagenomes is unusual because they are not commonly reported^8, 22^. This leads to the speculation that mangrove soil is a potential reservoir of pathogenic bacteria but further work is required to verify this finding.

In our RA soil metagenome, antibiotic and toxic compound resistance genes were also detected frequently. Their abundance may be related to the high percentages of Actinobacteria, which are known to produce a myriad of antibacterial compounds, and Deltaproteobacteria, members of which are known to be resistant to heavy metals and to oxidize heavy metals to their benign forms^23^, in our two tropical mangrove soil samples. We also obtained a high hit rate of the stress response gene in our soil metagenomic libraries, suggesting that the tropical mangrove environment is harsh for microorganisms. This is most probably due to polluted marine waters and high salinity and low aeration available in the muddy mangrove soil.

The biochemical tests showed considerably high amount of phosphorus and sulfur in our soil sample. However, the gene ontology analysis revealed that the genes associated with the metabolism of compounds containing phosphorus and sulfur were of relatively low abundance, suggesting that these compounds may mainly be involved in redox reactions in electron transport but not in microbial metabolism. This also implies that sulfur and phosphorus compounds exist in stable forms as mangroves are sink for inorganic compounds.

Although Actinobacteria and Firmicutes were two of the detected sub-dominant phyla, our analysis showed low frequency of spore producing bacteria such as *Bacillus* and *Streptomyces.* This may explain why the genes for sporulation and dormancy were not detected in our soil metagenomic DNA.

## Conclusion

This study demonstrated the high level of microbial diversity in mangrove swamps compared to the limited vegetation that is able to survive in this environment. The high abundance of members of Deltaproteobacteria and heavy metal and toxic compound resistance genes indicates that microorganisms have potential for bioremediation of heavy metals. The differences in the distribution of microorganisms compared with other previous studies on mangrove soils are most likely due to the different geographical locations. To the best of our knowledge, this is the first study of microbial diversity of mangrove soil in Malaysia using the NGS metagenomic approach. More mangrove soil samples collected from different locations in Malaysia are required to be analysed by this approach before a more defined conclusion on the microbiome of Malaysian mangrove soils and its functional genes can be reached.

## Acknowledgements

Kok-Gan Chan thanks the University of Malaya for the High Impact Research Grants (UM.C/625/1/HIR/MOHE/CHAN/01, Grant No. A-000001-50001 and UM-MOHE HIR Grant

UM.C/625/1/HIR/MOHE/CHAN/14/1, no. H-50001-A000027). There are no conflicts among authors regarding this research.

